# Computer-aided diagnosis of reflectance confocal images to differentiate between lentigo maligna (LM) and atypical intraepidermal melanocytic proliferation (AIMP)

**DOI:** 10.1101/2022.05.10.491423

**Authors:** Ankita Mandal, Siddhaant Priyam, Hsien Herbert Chan, Bruna Melhoranse Gouveia, Pascale Guitera, Yang Song, Matthew Arthur Barrington Baker, Fatemeh Vafaee

## Abstract

Lentigo maligna (LM), a form of melanoma in situ that predominantly affects sun-exposed areas such as the face, has an ill-defined clinical border and has a high rate of recurrence. Atypical Intraepidermal Melanocytic Proliferation (AIMP) is a term used to describe the melanocytic proliferation of an uncertain malignant potential. Clinically and histologically, AIMP can be difficult to distinguish from LM, and indeed AIMP may in some cases progress to LM. Reflectance Confocal Microscopy (RCM) is often used to investigate these lesions non-invasively, however, RCM is often not readily available nor is the associated expertise for RCM image interpretation. Here, we demonstrate machine learning architectures that can correctly classify lesions between LM and AIMP on stacks of RCM images. Overall, our methods showcase the potential for computer-aided diagnosis in dermatology, which in conjunction with the remote acquisition, can expand the range of diagnostic tools in the community.

## Introduction

Reflectance confocal microscopy (RCM) is an in vivo imaging modality that enables large cutaneous lesions in cosmetically sensitive areas to be visualised to the depth of the papillary dermis without the requirement of a biopsy for formal histological assessment. The changes seen in Lentigo maligna (LM) and atypical intraepidermal melanocytic proliferation (AIMP, elsewhere known as atypical melanocytic hyperplasia, or AMH) involve the levels above the papillary dermis and are thus ideal candidates for the use of RCM for diagnosis (Koller et al., 2009; Rocha et al., 2022).

Distinguishing between AIMP and LM is important as AIMP may only require ongoing monitoring while LM usually requires some form of definitive treatment before it may progress to invasion and possibility of metastasis (lentigo maligna melanoma). AIMP, in contrast to LM, can continue to be monitored *in vivo* and tends not to respond to topical or radiotherapy treatments (Rocha et al., 2022). Thus, correct diagnosis determines the level of treatment required. A number of clinical, histological, and reflectance confocal microscopy (RCM) criteria have been proposed and validated to assist in distinguishing AIMP and LM. RCM findings suggestive of LM include major criteria: non-edged papillae and round large pagetoid cells, and minor criteria: three or more atypical cells at the dermoepidermal junction in five RCM fields, follicular localisation of atypical cells, and nucleated cells within the dermal papillae. The presence of a broadened honeycomb is a significant negative feature for LM and is more suggestive of a benign seborrheic keratosis (Guitera JID 2010). Nevertheless, it can be difficult to distinguish early LM from AIMP given the common histological features of basal atypical melanocytic hyperplasia (Gómez-Martín et al., 2017). Further complicating the issue, AIMP has been shown to be, in fact, LM on further excision in 5% of cases (Bou-Prieto et al., 2021). Predictors of AIMP progression to LM have not been well defined, though could include a target-like pattern and high-density vascular network on dermoscopy, and the presence of contact between dendritic cells on RCM (Rocha et al., 2022). This may indicate that a binary classification approach to diagnosis is a simplification of an underlying spectrum of pathologies between AIMP and LM.

RCM enables longitudinal study of large heterogeneous lesions, tracking heterogeneity spatially and change over time non-invasively over multiple follow-ups and thus, may be a more suitable modality for investigation of AIMP and LM. Despite a treatment code, which acknowledges the utility of RCM based diagnosis, access to RCM and specialist interpretation of RCM images is often a limitation to its use, and thus, there is a need for automated approaches to improve medical equity. For example, in Australia, the country with the highest prevalence of LM, only ~10 RCM exist with a similar number of clinicians trained to interpret such images. Furthermore, a gold standard for borderline or uncertain malignancy does not exist and current criteria are neither reproducible nor accurate (Elmore et al., 2017). Computer-aided diagnosis can help address the issue of access to diagnostics since the diagnosis and image acquisition can be physically separated (remote acquisition), and either entirely computational diagnosis or computer-aided diagnosis by the clinician can allow far greater patient throughput. Computer-aided diagnosis has further applications for the prediction of prognosis, and machine-learning has been employed in prostate and breast cancer to determine grades of differentiation which hold clearly defined risks of progression and prognostic outcomes (Fusano et al., 2020; Khan et al., 2021).

Machine learning approaches in dermatology to date have typically focused on the distinction between benign and malignant lesions based on clinical images (Aractingi and Pellacani, 2019; Pérez et al., 2021; Petrie et al., 2019). For this, extensive public libraries exist and dermatologist-level performance has been achieved with a CNN pre-trained on a fine-tuned subset of 130,000 images derived from the ImageNet dataset (Esteva et al., 2017). A similar approach to RCM images has been hampered by the limited availability of RCM infrastructure and labelled datasets, and the requirement of extensive pre-processing, segmentation and feature extraction prior to classification.

While RCM can image, with a cellular resolution, at a range of depths to reconstruct a 3D volume for an associated tissue, the classification of these 3D volumes is less established than the classification of 2D images using computer vision approaches. Processing of 3D volumes is also much more computationally expensive than 2D image analysis. One way to address this is to project 3D information into 2D, which can then be further analysed using various types of machine learning approaches. In this work, we have chosen to leverage the LZP projection, which is the latest fast approaches for projecting a 3D image into 2D while preserving information (Haertter et al., 2021).

Successful machine learning RCM decision support systems have been employed in the diagnosis of BCC (Campanella et al., 2021) with comparable performance achieved with a deep-learning based model and with RCM experts. Other applications have been in the diagnosis of congenital pigmented macules in infants (Soenen et al., 2021). More extensively, deep neural networks have been employed in RCM image quality assessment, assisting the interpretation of RCM mosaics, and automated detection of cellular and architectural structures within the skin (Bozkurt et al., 2021; D’Alonzo et al., 2021; Kose et al., 2020, 2021; Wodzinski et al., 2020).

Here, our hypothesis was that the projection of 3D virtual stacks into single 2D images could deliver high accuracy machine binary classification between LM and AIMP lesions with lesser computational requirements, primarily memory. Our aim was to demonstrate high accuracy machine classification of LM and AIMP lesions utilising projections of RCM stacks that had been validated by clinician diagnosis and subsequent biopsy. Our approach is fast and implementable on minimal computational architectures and achieves high accuracy to showcase the potential for computer-aided diagnosis for these pathologies.

## Methods

### Study design and participants

The study population comprised a total of 151 patients, who attended the Sydney Melanoma Diagnostic Centre (Camperdown, NSW) RCM clinic between January 2019-December 2020 who had biopsy-proven LM or AIMP lesions. RCM stacks were obtained for these patients from the RCM image database (HREC/11/RPAH/123 - X15-0392 Sydney Local Health District Ethics Review Committee (RPAH zone)).

### RCM acquisition procedure and exclusion procedure

Clinically identified atypical pigmented lesions were scanned using a handheld Vivascope 3000 (Vivascope ID). Areas representing the diagnosis were identified by a trained confocal specialist, and stacks of 32 images were sampled at each site. Stacks were excluded when they were targeted at margins of the lesions. Following imaging, areas with RCM-detected atypia were biopsied, and pathology was confirmed via formal histological diagnosis to create our ground truth. For slice-level classification, the clinician revisited each stack and, for each individual image in the stack, assigned a diagnosis of LM, AIMP or neither.

### Image processing

Individual images were exported from microscope software Vivascan (Vivascope ID) as 24-bit TIFF single images according to z-slice. Folders of individual TIFFs were imported into FIJI (ImageJ reference) as a virtual stack, and then initial projections were calculated using z-projection with the maximum and median. For subsequent classification using predictive modelling, stacks were projected using the FIJI plugin for local z projection (LZP) (https://biii.eu/local-z-projector), an optimal method for structure-specific projections that can be computed rapidly (Haertter et al., 2021). LZP was run in default settings for these stacks for the reference surface, with a max of the mean method with a 21 pixel neighbourhood search size and a 41 pixel median post-filter size, and using MIP to extract the projection. Projections were then exported as 8-bit JPGs (1000 × 1000 pixels) and uploaded to Google Drive where they were read using cv2.imread (Bradski, 2000) and resized to 256 × 256 pixel images. Augmentation was performed only on the AIMP data set using cv2 similarly (8 images augmented to 32 images by adding either horizontal flip or vertical flip or both horizontal and vertical flips). Resizing to 256×256 pixels was done using cv2 resize function with inter-cubic interpolation. For the slice level ternary classification, individual TIFFs were read in using cv2.imread and resized to 256 × 256 pixel images.

### Predictive Modelling

#### Model development

Different popular convolutional neural network (CNN) architectures were employed to classify AIMP vs LM projections, including ResNet50 (He et al., 2015), ResNet101 (He et al., 2016), InceptionV3 (Szegedy et al., 2016), VGG16 (Simonyan and Zisserman, 2015), and DensNet169 (Huang et al., 2018). These models were pre-trained on ImageNet (https://www.image-net.org), and the model parameters were then fine-tuned on our projections of RCM stacks. We also developed a 6-layer CNN to evaluate the predictive performance on a simple architecture that is potentially less prone to overfitting. The Adam optimisation algorithm (Kingma and Ba, 2017) was adopted to optimise the learning rate of neural network parameters for all the architectures except for ResNet50 and InceptionV3, for which the RMSProp algorithm (Kurbiel and Khaleghian, 2017) was used. Images were augmented to increase sample sizes. The strategy used for augmentation was flipping (vertical, horizontal, and a combination of both). The performance of the models was assessed on the *validation* set, a subset of projections that was held back from the training projections and used to give an estimate of model’s accuracy while tuning model’s parameters. As detailed in the next subsection, the best-performing model as per the validation accuracy was further evaluated on a completely unseen *test* set containing projections not accessible during the model training to mitigate overfitting risk and select a model with more a generalisable performance.

To extend the diversity of the models evaluated, we also combined deep-learning-based feature extraction with other traditional classifiers. Accordingly, latent features were extracted from the DenseNet169 and ResNet50 models (i.e., the first and second best-performing CNN models). Extracted latent features derived from ResNet50 have shown better performance once used as predictive variables of different commonly-used classifiers, including support vector machines (SVM), random forest (RF), and k-nearest neighbours (KNN), and AdaBoost (Wang, 2012). Models were developed in Python using Keras neural network library on the TensorFlow platform. For slice level classification, an additional activation function, SoftMax, was included in order to perform ternary classification.

#### Model validation and performance metrics

The *k-*fold cross-validation (Kohavi, 1995) was employed for model validation to give a more robust and generalisable estimate of the model’s predictive performance. Accordingly, patients (not images) were split into test and train sets. The test set was held out, and the training set was randomly partitioned into *k* complementary subsets; one is taken as a validation set for model optimisation and the rest as the training set. Projected images were randomly split into test and train sets with a constraint that multiple projected stacks from a single patient were included in either test or train sets (i.e., patient-level splitting) to avoid any potential information linkage from train to test set. Accordingly, roughly 20% of projections were withheld as a test set. This process was repeated *k* times so that each subset would be considered as a validation set in one iteration. The performance metrics over the holdout test set were then evaluated and reported for each of the *k* models trained. We performed a 5-fold crossvalidation, and in each iteration, we used multiple metrics to measure the prediction performance on the test set, including accuracy (rate of correct classifications), recall or sensitivity (true positive rate), precision (positive predictive value), and F1-score that is the harmonic mean of the precision and recall, i.e., F1-score = 2/(recall^-1^ + precision^-1^). The quality of models was also depicted by the receiver operating characteristic (ROC) curve, which plots the true positive rate (i.e., sensitivity) against the false positive rate (i.e., 1-specificity) at various threshold settings (Hoo et al., 2017). The area under the ROC curve (AUC) was computed, which varies between 0.5 and 1. The higher the AUC, the better the performance of the model at distinguishing between AIMP versus LM; a random or uninformative classifier yields AUC=0.5. The confusion matrix was also reported on the selected model detailing the total number of correct and incorrect predictions – i.e., true positives (TP), false positives (FP), true negatives (TN), and false negatives (FN). For a sensible model, the diagonal element values will be high (TP and TN), and the off-diagonal element values will be low (FP and FN).

### Prediction Interpretation

We used Gradient-weighted Class Activation Mapping (Grad-CAM) (Selvaraju et al., 2017) [2] algorithm to produce visual explanation heatmaps highlighting the important regions in the images that contribute to the decision made by the best-performing CNN model (i.e., DenseNet169). Accordingly, AIMP and LM projected images in the test sets were run through the DenseNet169 model that is cut off at the layer for which we want to create a Grad-CAM heatmap. The layer output and the loss were then taken, and the gradient of the output of the model layer with respect to the model loss was found. The gradient which contributes to the prediction was taken, reduced, resized, and rescaled so that the heatmap can be overlaid with the original image. The implementation was made in Python using the TensorFlow platform and is available in the study’s code repository (*c.f*., Code and Data Availability).

### Statistical analysis

The statistical hypothesis tests comparing the significance of the performance enhancement comparing the best performing method (DenseNet169) and other competing algorithms were conducted using the paired two-tailed t-test. Statistical significance was defined as a *p*-value < 0.05. Statistical analyses were performed in R using the ‘stats’ library.

## Results

The information of the patients is detailed in **Supplementary Table 1**. Overall, 541 RCM stacks of 28 – 40 images (750 μm – 750 μm with 3.5 – 5.0 μm depth spacing) were collected from 135 patients. **Figure 1A** illustrates the image processing and diagnostic modelling pipeline developed in this study. The imbalance in the proportion of LM versus AIMP cases was partially handled via augmenting AIMP images by flipping them horizontally, vertically, and in both directions. Together, the training set included 537 projections (389 labelled LM and 148 AIMP) and the test set comprised 115 projections (83 LM and 32 AIMP).

**Figure 1.**
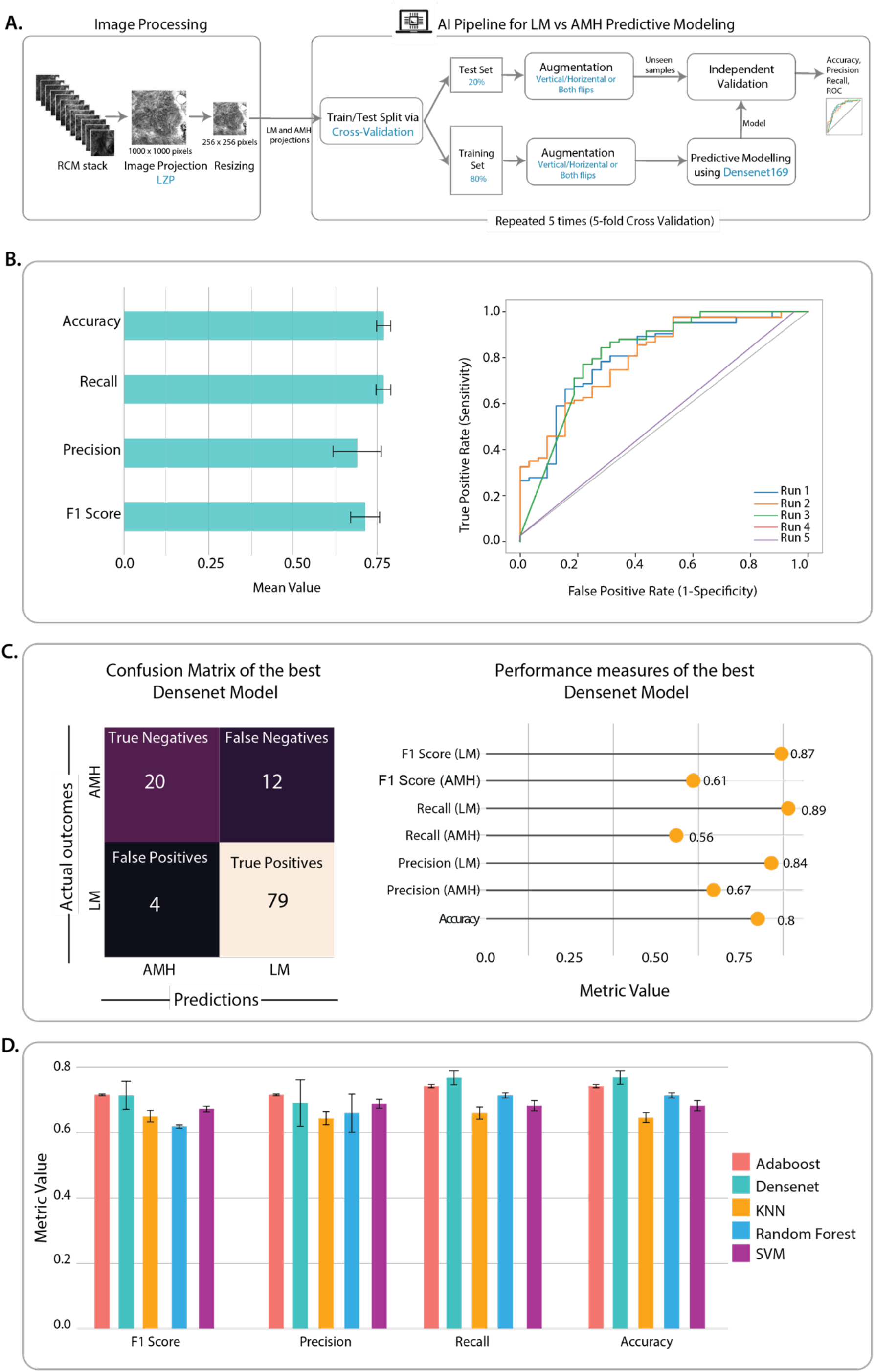
**A.** The schematic workflow of the study comprising image processing (projection and resizing), and deep learning model development and validation. **B.** The test-set performance of DenseNet196 model over five runs of cross-validation is represented as bar plots and receiver operator characteristic (ROC) curves. The bar plots represent the weighted-average of the performance metrics (accuracy, recall, precision, and F1-score) across five runs. The error bar represents the standard error. **C.** The confusion matrix representing the details of predictions made by the best-performing DensNetl96 model (Run 1) and performance metrics in predicting LM and AIMP projections in the corresponding test set (20% of held-out data in Run 1 of 5-fold cross-validation). D. The Comparison of the DenseNet196 classifier with the traditional machine learning algorithms (AdaBoost, k-nearest neighbour (KNN), Random Forest, and Support Vector Machine (SVM)); the bar plots represent the weighted-average of the performance metrics (accuracy, recall, precision, and F1-score) across five runs. The error bar represents the standard error.

Among selected CNN architectures pre-trained on the ImageNet dataset, DenseNet169 achieved the highest predictive power on the validation set (*validation* accuracy = 0.84). The predictive power of DenseNet169 was assessed on the test set (115 unseen images) using multiple metrics, including accuracy, weighted-average recall, weighted-average precision, and F1-scores for models developed via cross-validation (**Figure 1B**). The class-specific precision and recall were averaged with the consideration of the class imbalance (i.e., weighted average). The best-performing DenseNet169 model was achieved via the first run of cross-validation (*c.f.* Run 1 in Figure 1B, ROC curves) with the accuracy of 0.80 on the test set (**Figure 1C**). The *test* accuracy of DenseNet169 as a standalone feature learning and classifier was higher than traditional classifiers (using default hyper-parameters) (**Figure 1D**). However, the performance improvement was only significantly higher as compared to SVM and KNN (*p*-value < 0.05, paired, two-tailed t-test). Since DenseNet169 performed better or on par with the other classifiers, it was used for the subsequent patient-level prediction interpretation.

We examined predictions made by DenseNet169 models for each of 115 projected images in the test set across five runs of cross-validation (**Figure 2A**). Image IDs in this figure can be mapped to the corresponding RCM stacks using **Supplementary Table 2**. To further understand factors contributing to the model’s false or true predictions, we plotted Grad-CAM heatmaps of selected images (**Figure 2B**) from the test set. The selection criteria were to include examples of LM and AIMP patients that are correctly classified (i.e., a true positive and a true negative) as well as examples of incorrectly diagnosed images (i.e., a false positive and a false negative) across the majority of the runs. We limited the selection to non-augmented images. The Grad-CAM heatmaps of the remaining test images are available in the GitHub repository (see Code Availability).

**Figure 2.**
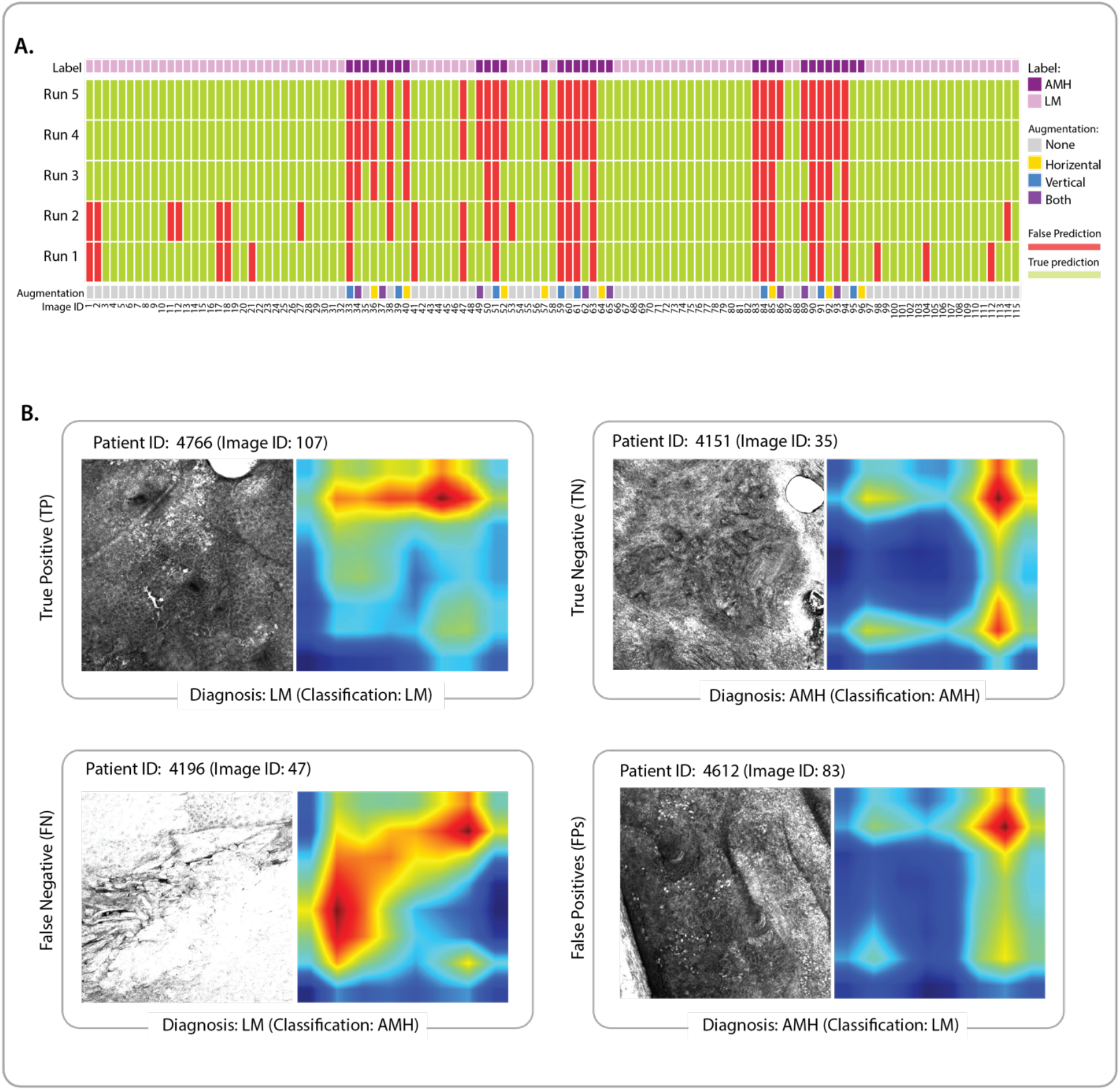
**A.** Patient-level predictions of LM and AIMP images in the test set across five runs of the cross-validation. The heatmap represents the false predictions (false positives and false negatives) in red and correct predictions (true positives and true negatives) in light green. Each 2D projection image (equivalent to an RCM stack is identified by a unique ID (Supplementary Table 1) and colour-coded based on the diagnosis (LM or AIMP) and augmentation of the 2D projections (vertical flip, horizontal flip, both, and none, i.e., no augmentation). **B.** Selected projections in the test set and their corresponding Grad-CAM heatmaps enabling the interpretation of false and true predictions of LM (positive) and AIMP (negative) diagnoses.

To examine the effect of the projection, we visually compared projections using LZP with slice-by-slice clinician diagnosis to examine how well LZP projection preserved diagnostic markers in our original RCM stacks. In general, while diagnostic markers were not recognisable by the clinician in the projected image, the classification was still accurate in that image information was preserved, and classification could be successful. Representative images are shown for each class in Fig. 3, alongside maximum z-projection (the highest pixel intensity at each location) and the median z-projection (the median pixel intensity at each location). Figure 3A indicates a representative True Positive, that is, an LM-diagnosis classified as LM where the stack had atypical enlarged melanocytes and dendritic cells present at superficial levels indicating pagetoid spread. This was preserved in the projection, indicating that melanocytes were present at most levels within the stack. Figure 3B shows a representative True Negative, that is, an AIMP-diagnosis that is classified as AIMP. The stack showed diffuse enlarged melanocytes at the basal layer with no dendritic cells. In the projection, the air bubble artifact in the top right is preserved, though did not interfere with the correct classification being made. Figure 3C shows a representative false positive, that is, an AIMP-diagnosis classified as LM. There the stack had diffuse enlarged melanoyctes at basal layer, with no pagetoid spread and no dendritic cells. The melanocytes were retained by projection. However, the information regarding at which depth the melanocytes were located was removed during projection. Lastly, Figure 3D shows a representative False Positive, that is, an LM-diagnosis classified as AIMP. There the stack was acquired too early in superficial skin layers, and the presence of a skin fold prevented the acquisition of the whole en-face image. Pagetoid spread of non-dendritic melanocytes was present; however, irregular skin surface and non-perpendicular z images made it hard to interpret pagetoid spread.

**Figure 3.**
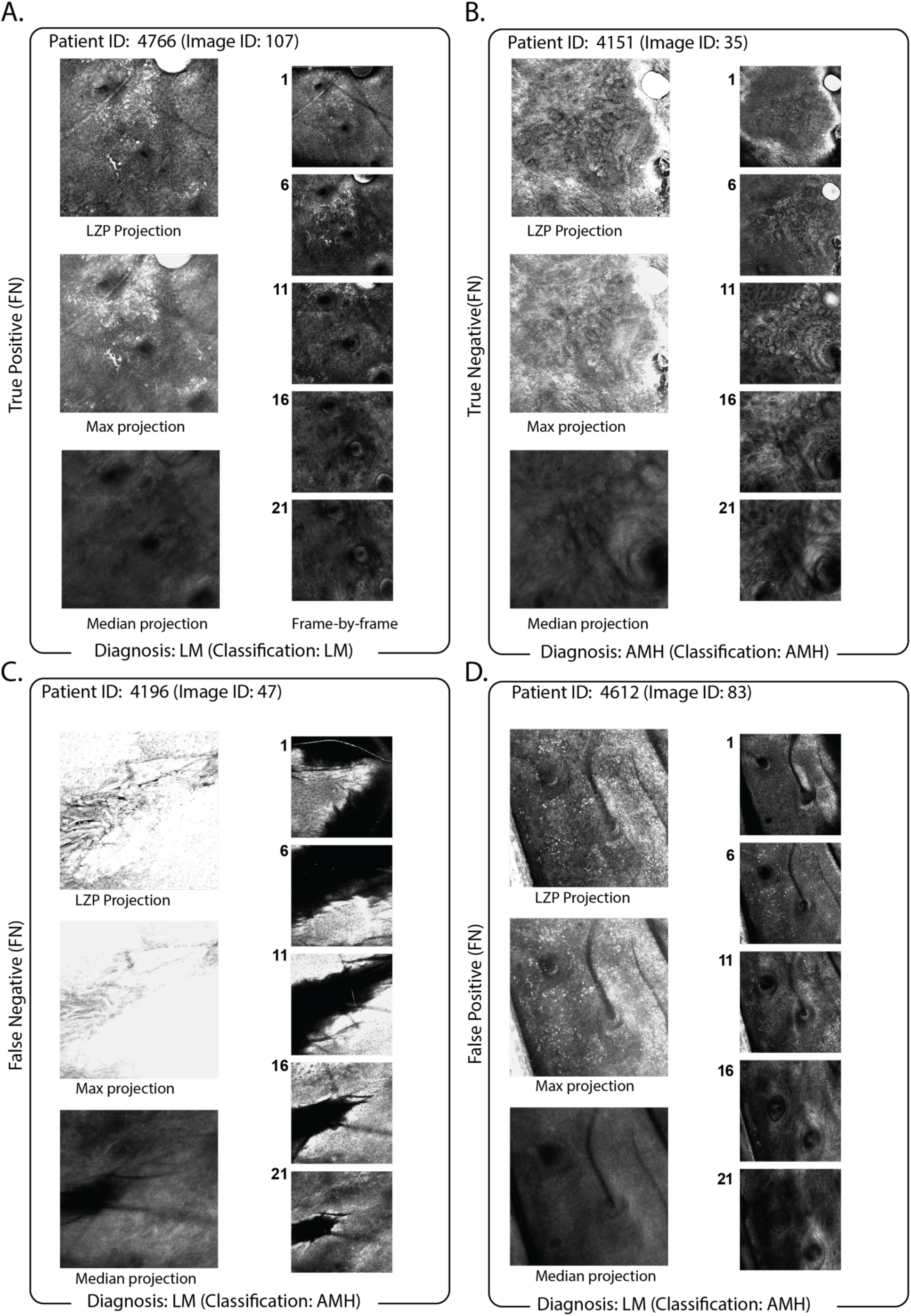
Comparison of LZP projection vs max- and median-projection for exemplary classification outcomes. Exemplary data for (A) LM-diagnosed image stack correctly classified at L; (B) AIMP-diagnosed image stack classified as AIMP; (C) LM-diagnosed image stack misclassified as AIMP; and (D) LM-diagnosed image stack misclassified as LM. For all panels, projections are shown on left (LZP: top; max-projection: middle; median-projection bottom) with individual slices at specific depths (z = 1, 6, 11, 16, 21) shown inset on right. Max-projection is generated by taking the maximum value pixel across all slices of the stack, median-projection is generated by taking the median value pixel across all slices of the stack.

LZP can be seen to outperform simple max-projection since the individual detail and diagnostic markers remain clear (e.g. Figure 3A, Figure 3B true positive and true negative, respectively). However, when there are individual frames that are saturated at maximum brightness, these can dominate the signal in the projection (Fig. 3C), and where the image stack is bright in different regions, this local information is lost upon projection. Likewise, in Fig 3D, marker information that shows clearly enlarged melanocytes at the basal layer (Fig. 3 inset) is potentially misinterpreted as being present at all slices of the stack when considering only the projection.

We compared the classification of projections to the classification of individual slices at the slice by slice level. We revisited all stacks to add clinician diagnosis to individual slices as containing LM features, AIMP features or non-pathological skin layers, respectively since not all slices in a stack contained pathology. This increased the total number of images but also altered the problem to a ternary classification problem. The best performing models for this ternary classifier were SVM and KNN, with average test accuracies of 0.59 and 0.70, respectively (with Resnet101 used for feature extraction). However, it is critical to note that these average accuracies included entire classes that were not classified correctly. For example, for AIMP, the recall was zero; that is, no slices diagnosed as AIMP were correctly classified as AIMP using this slice level ternary classifier.

## Discussion

Machine learning approaches to RCM analysis strive to enable more efficient image interpretation by a human user and also to assist with the classification of skin lesions. In general, RCM image analysis is hindered by artefacts both within an individual field of view and in mosaics where multiple RCM fields are stitched together. Differences in exposure and image intensity can also make mosaic rendering difficult. The dermoepidermal junction (DEJ) is where specific pathology is located in a number of diagnoses, particularly in AIMP and LM, and thus automated identification of the DEJ location and relevant cellular structures can help to focus attention to areas of a stack that may offer a high diagnostic yield. Machine learning-driven processes to assess the quality of RCM mosaics and identify the DEJ and other cellular structures have been effective in improving the workflow of RCM image interpretation (D’Alonzo et al., 2021, 2021; Kaur et al., 2016; Kose et al., 2020, 2021).

Machine learning approaches specifically to the classification of RCM images of skin lesions have focused until now on BCC and melanoma diagnosis (Campanella et al., 2021; Wodzinski et al., 2020), lentigos (Halimi et al., 2017a, 2017b; Zorgui et al., 2020), and congenital pigmented macules (Soenen et al., 2021).

The main features used to differentiate between LM and AIMP both histologically and on RCM are the confluence and location of atypical melanocytes (Rocha et al., 2022). In LM, these are present at all levels of the epidermis; however, in AIMP, whilst the melanocytes are enlarged, there is no pagetoid spread, and they are present only in basal layers. However, the biological differentiation between AIMP and LM is not entirely clear cut (Ensslin et al., 2018). A decision support system in this context can thus not only assist in diagnosis but may also identify specific features that may differentiate the two conditions and hence provide insight into the biological nature of AIMP vs LM.

We optimised our model to deliver a binary classification that could differentiate between AIMP and LM samples with a test accuracy of 0.80. Our approach was robust in that we were agnostic to a particular architecture, trying a variety of approaches and testing which had the highest accuracy and AUC. For different pathologies or diseases, a similar agnostic approach could be applied to the dataset to identify the architecture best suited for efficient and accurate classification and diagnosis.

Training data sets for these previous studies have included single images sometimes pre-selected in the vicinity of the DEJ (Soenen et al., 2021), RCM mosaics (Wodzinski et al., 2020) or 3D reconstructions (Zorgui et al., 2020). In contrast, we utilised a projection approach to project 3D and volumetric image data into a 2D representation of that volume as it offered some specific benefits. First, our computational performance was significantly optimised since we did not have to run large image stacks of raw TIFF image data. We could instead use compressed single JPG images, greatly reducing the memory overhead (projection ~ 500 kB; stack ~ 100 MB). If the projection method is optimised to preserve diagnostic markers, then diagnostic markers of relevance that can be shared over many layers, but not all, of the volume should be emphasised in the final projection, and thus a classifier can utilise these to make its prediction.

Projection is of course, not without drawbacks. First, it requires good alignment between the individual slices of a stack, and it is influenced by any drift in x- and y- as the operator moves deeper into the tissues. Where there is large drift, the separate slices are not registered relative to each other and the projection will blur features during projection and may result in loss of diagnostic markers. Similarly, where individual slices are saturated or overly bright, this saturated signal may dominate in the final projection. An example of this is shown in Fig. 3C where saturation in individual slices is localised to specific regions, but upon projection, this data is preserved, and so the entire image is saturated, in that instance resulting in misclassification.

The alternative to projection is to run a slice-by-slice classifier. However, this requires a clinician to provide a slice-by-slice diagnosis at the slice level, which is time-consuming. Furthermore, it necessitates a ternary rather than binary classifier since there will be many slices that, in fact, contain no specific features for LM or AIMP. Thus, to test a slice-by-slice classifier appropriately, a ternary classifier must be used, which is even more susceptible to class imbalance. Our best accuracies achieved via ternary classification at the slice level were 0.70, with total misclassification of our AIMP cohort.

RCM-trained clinicians typically train on individual slices in the overall volume and make their diagnosis while imaging through the disease tissue. They typically start the imaging above the disease tissue and continue through the tissue to image past it. This extra information can confound machine classification, especially when an artefact is present, such as an air bubble or a follicle, as this information will be more prominent in the final projection. Clinicians make their diagnostic assessment from cellular appearance and not from a projection. In projections, depending on the approach used, local bright detail will be emphasised, and non-disease tissue may appear as disease tissue. Clinicians/ technicians could adapt their imaging approach in order to derive more benefit from computer-aided diagnosis in the future by avoiding drift in x- and y-, not projecting past the pathology or imaging too early, and avoiding saturation in any slice of the overall stack (adjusting the exposure, laser intensity, or the imaging conditions to guard against this).

The rarity of RCM instrumentation and the paucity of skills and expertise with these instruments, particularly in remote areas with a high prevalence of melanoma, indicates that remote and computer-based diagnosis has much to offer (Rao? JAAD 2013). Reduction in diagnostic time has been achieved with ML-driven pre-selection of specific images that need expert review in the context of prostate cancer(Campanella et al., 2021). A similar approach may be achieved with the method illustrated in this paper. Microscopy technology is improving rapidly and will continue to miniaturise further; as such, it may be much easier to locate an RCM or low-resolution microscope to a patient than to bring the patient and the expert together. Further training of machine learning classifiers, as well as training of operators in preparing ‘machine-friendly’ image stacks, will benefit patient outcomes in the field and the further implementation of computer diagnosis as technologies improve.

## Abbreviations

LM: lentigo maligna
AIMP: Atypical Intraepidermal Melanocytic Proliferation
RCM: reflectance confocal microscopy
LZP: local z projection

## Data and Code Availability

All RCM images and codes are available for non-commercial uses at https://github.com/VafaeeLab

